# Reproduction costs can drive the evolution of groups

**DOI:** 10.1101/325670

**Authors:** Yuriy Pichugin, Arne Traulsen

**Affiliations:** Max Planck Institute for Evolutionary Biology, August-Thienemann-Str. 2, 24306 Plön, Germany

## Abstract

A fascinating wealth of life cycles is observed in biology, from unicellularity to the concerted fragmentation of multi-cellular units. However, the understanding of factors driving the evolution of life cycles is still limited. We investigate how reproduction costs influence this process. We consider a basic model of a group structured population of undifferentiated cells, where groups reproduce by fragmentation. Fragmentation events are associated with a cost expressed by either a fragmentation delay, a fragmentation risk, or a fragmentation loss. The introduction of such fragmentation costs vastly increases the set of potentially optimal life cycles. Based on these findings, we suggest that the evolution of life cycles and the splitting into multiple offspring can be directly associated with the fragmentation cost. Moreover, the impact of this cost alone is strong enough to drive the emergence of multicellular groups, even under scenarios that strongly disfavour groups compared to solitary individuals.

## 1 Introduction

All living and evolving organisms are born, grow and reproduce, giving birth to new organisms [van Gestel and Tarnita, 2017, Stearns, 1992, Maynard Smith and Szathmáry, 1995, Bonner, 1998, Roze and Michod, 2001, Pfeiffer and Bonhoeffer, 2003, Rainey and Kerr, 2010, Ratcliff et al., 2012, Hammerschmidt et al., 2014, De Monte and Rainey, 2014, Kaveh et al., 2016]. Natural selection promotes those organisms that perform this cycle in a more efficient way than others, as these produce more offspring per time. Surprisingly, even the simplest organisms demonstrate a great variety of reproduction modes: *Staphylococcus aureus* produces independent propagule cells [Koyama et al., 1977], cyanobacteria filaments fragment into multicellular threads [Rippka et al., 1979] while *Gonium pectorale* disperses into independent cells [Stein, 1958]. These instances show that there is no universally optimal reproduction mode. Instead, the way how cell groups produce offspring is an adaptation to the environmental conditions and constrained by the biological properties of the organism [van Gestel and Tarnita, 2017].

One such property which can limit the possible life cycles is the group fragmentation cost. There is substantial evidence that reproduction is costly in natural populations. For example, during the fragmentation of a simple multicellular organisms, the release of cells requires the break of the cell matrix, which takes time and resources [Birkendal-Hansen, 1995, Basbaum and Zena, 1996]. Also, not every cell may pass to the next generation of groups, for instance in slime molds cells forming the stalk of the colony die shortly after the spores are released [Bonner, 1959]. Another example are cells constituting the outer layer of a *Volvox carteri* colonies - these cells die upon the colony reproduction [Smith, 1944]. Combined, this evidence shows that reproduction can be associated with a conspicuous cost.

There are only a few studies of the evolution of reproductive modes which explicitly take into account the fragmentation cost. Libby et al. [2014] modelled the evolution of life cycles of colonial forms of *Saccharomyces cerevisiae*. In their model, the fragmentation of tree-structured cell clusters was attributed to the death of cells. These cells become weak links and loose connections with neighbouring cells causing fragmentation of the cluster. However, while Libby et al. considered a detailed model of binary fragmentations of cell clusters, they did not investigate the whole range of fragmentation outcomes. In previous work, we have extensively analysed all possible ways of group fragmentation and found evolutionary optimal life cycles under various fitness landscapes [Pichugin et al., 2017]. For costless group reproduction, only binary fragmentation, where a larger group splits into two parts, can be evolutionary optimal in terms of maximising population growth. The same holds for the case of proportional cost, where upon division into s parts, *s* − 1 cells die. However, for fragmentation with a fixed cost in a form of a single cell loss, fragmentation modes with multiple offspring can become evolutionary optimal.

In this study, we investigate the influence of the fragmentation cost on the evolution of “staying together” life cycles [Tarnita et al., 2013]. We explicitly incorporate fragmentation costs arising from three scenarios: fragmentation delay, fragmentation risk and cell loss. We discuss the set of life cycles which can be evolutionary optimal for costly fragmentation. Then, we investigate how the distribution of optimal life cycles on a set of random fitness landscapes depends on the value of the fragmentation cost. Finally, we consider in detail those fitness landscapes in which the increase in a group size always reduces the performance of the group, i.e. the fastest growth and the best protection is achieved by independent cells. We show that even in these fitness landscapes that strongly disfavour multicellular groups, fragmentation costs can promote the evolution of life cycles involving the emergence of multicellular groups.

## 2 Methods

### 2.1 Growth and death of groups

We consider a population composed of unstructured groups (or complexes) of cells, which emerge, grow and fragment into offspring groups, thus completing the life cycle. Groups grow by dividing cells staying together after reproduction [Tarnita et al., 2013]. Due to the absence of any structure, the properties of a group are determined by its size *i* alone. We denote the abundance of groups of *i* cells in a population as *x*_*i*_. We additionally assume that the size of groups in a population is bounded by *n*. Groups of size *i* have a death rate *d*_*i*_ and cells in a group have the division rate *b*_*i*_, thus the growth rate of a group is *ib*_*i*_. The vectors of birth rates **b** = (*b*_1_,…, *b*_*n*_) and of death rates **d** = (*d*_1_,…, *b*_*n*_) define the fitness landscape of the model, see Fig. 1a.

### 2.2 Group fragmentation

New groups are produced by the fragmentation of existing groups. We further assume that the fragmentation occurs immediately after the growth of the group. Thus, upon each cell division, a group grows in size by one and either remains in this state until the next cell division, or splits into two or more smaller groups. As any group can be characterized by the number of cells comprising it, any fragmentation or growth can be characterized by a partition of this integer number. A partition is a way of decomposing an integer *m* into a sum of integers without regard to order, summands are called parts [Andrews, 1998]. We use the notation *κ* ⊢ *m* to indicate that *κ* is a partition of *m*, for example 2 + 2 ⊢ 4, see Fig. 1b. The number of partitions of *m* grows fast with *m*. In the current study, we use *n* = 19 and thus *m* does not exceed 20. For *m* = 20, there are in total 2693 non-trivial partitions (with more than one part).

As example of using partitions to characterize fragmentation modes, consider a group of 2 cells in which the 3rd cell is born. If the group fragments without any cell dying, the product is either three independent cells (partition 1 + 1 + 1 ⊢ 3) or a group of two cells and an independent cell (partition 2 + 1 ⊢ 3). If a cell is lost upon fragmentation, the only possible result is two independent cells (partition 1 + 1 ⊢ 2). In the absence of fragmentation, the product is the single group of three cells (the trivial partition 3 ⊢ 3).

### 2.3 Three way of implementing fragmentation costs

We consider three qualitatively different scenarios that capture the fragmentation cost: fragmentation delay, fragmentation risk, and fragmentation loss.

**Figure 1:**
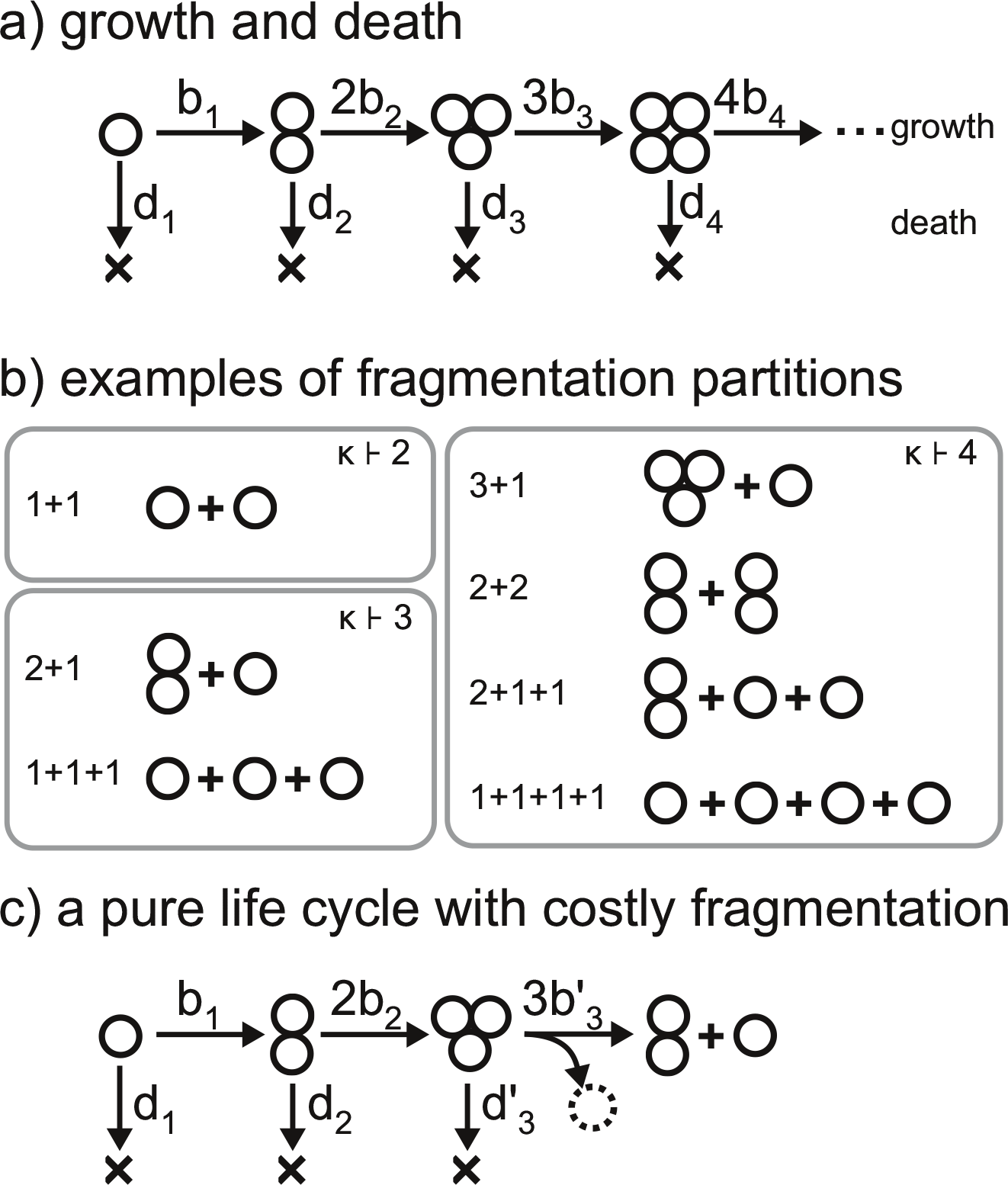
Model of life cycles. *(a)* The fitness landscape is defined by vectors of growth and death rates. Cells in a group of size *i* grow at rate *b*_*i*_ and groups die at rate *d*_*i*_. *(b)* The fragmentation of groups is described by a partition of an integer number into a sum of integers. All possible fragmentations of groups of size 2,3, and 4 are presented here. *(c)* In a deterministic life cycle, all groups follow the same partition at the fragmentation. For costly fragmentation, the growth rate at the maturity size may be smaller than prescribed by the fitness landscape 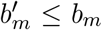, the death rate at the maturity size may be larger than prescribed by the fitness landscape 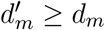 and some cells may be lost upon the fragmentation (one cell in the illustrated case).

#### 2.3.1 Fragmentation delay

In the case of the fragmentation delay, the process of fragmentation is not immediate and takes time *T*. This scenario covers situations where the fragmentation of the group requires the investment of resources, which otherwise would be spent on the further growth of the group. The transition time is inverse to the transition rate, thus we define the rate of fragmentation of a clusters of size *m* by

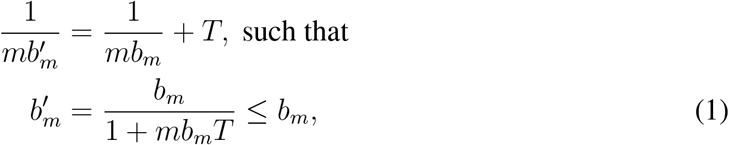

where *T* it the fragmentation delay. Consequently, this scenario can be captured by changing the fitness landscape in terms of the birth rate at the size prior to fragmentation.

#### 2.3.2 Fragmentation with risk of death

In the case of the fragmentation with risk, the organism expresses risky behavior prior to the fragmentation. For example, an organism could leave the shelter or break its shell in order to reproduce. Under this scenario, the risky behaviour increases the death rate at the final stage of the organism life cycle by *R*

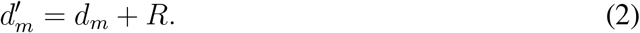

Again, this scenario corresponds to a change of the fitness landscape.

#### 2.3.3 Fragmentation with loss

For fragmentation with loss, *L* cells die as upon the group fragmentation, thus the combined size of offspring groups is by *L* smaller than the size of the fragmented cell cluster. Under this scenario, the fragmentation followed by the growth from size *m* to *m* + 1 is characterized by a partition *κ* ⊢ *m* +1 − *L*. We assume *L* to be constant, i.e. clusters loose the same number of cells independently on the partition of the parent group into offspring.

The three considered scenarios are not mutually exclusive, all three types of cost may be present simultaneously. However, for simplicity of the presentation of results, we illustrate each scenario of the fragmentation cost independently.

### 2.4 Population dynamics under a deterministic life cycle

For costless fragmentation, natural selection favours a narrow subset of life cycles, called deterministic life cycles in [Pichugin et al., 2017], see Fig. 1c. In these life cycles, groups always grow up to some maturity size *m* ≤ *n*, always fragment immediately after the *m* + 1-st cell is born, and the fragmentation always follow the same pattern, given by a single partition. Also for costly fragmentation, natural selection promotes only deterministic life cycles, see Appendix A.1. Thus, here we do not consider any life cycles other than deterministic ones, where a life cycle would follow several paths, sometimes fragmenting in one way and sometimes in another one.

Under a given deterministic life cycle, the state of a population can be described by abundances of groups *x*_*i*_ of each possible size *i* from one cell to *m* cells given by the vector (*x*_1_, *x*_2_, · · ·, *x*_*m*_). There are no groups of size *m* +1 or larger, because under deterministic life cycle, any group fragments immediately after the next cell is born in a group of the maturity size *m*.

The dynamics of the population state can be expressed in a form of the system of *m* differential equations: one equation for each particular size of groups. The change in the number of groups of a given size is influenced by growth, death and fragmentation. This leads to the set of equations

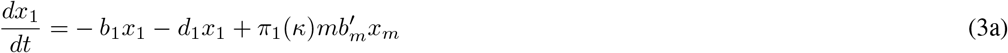

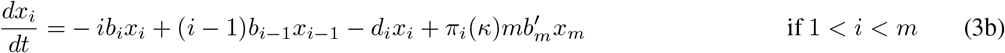

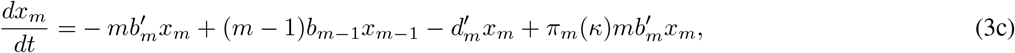

Here, Eqs. (3a) and (3b) describe the dynamics of the abundances of groups *x*_*i*_ that grow without fragmentation, because they do not reach the maturity size *m*. The first two terms in Eq. (3b) −*ib*_*i*_*x*_*i*_ + (*i* − 1)*b*_*i*−1_*x*_*i*−1_ describe the change in *x*_*i*_ due to the group growth. The next term −*d*_*i*_*x*_*i*_ describes the death of groups. The last term 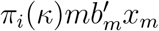 describes the emergence of new groups of size *i* resulting from the fragmentation of mature groups. The integer *π*_*i*_(*κ*) is the number of groups of size *i* that emerge in a single act of fragmentation according to the partition *κ*, and 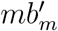 is the growth rate prior to fragmentation (see Eq. (1)).

Eq. (3c) describes the dynamics of groups of maturity size *m*, which will inevitably fragment according to the partition *κ* upon the next cell division. For fragmentation with delay, the rate of transition to the next state (fragmentation) is smaller than the cell birth rate 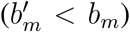 implied by the fitness landscape birth vector b (see Eq. (1)). For fragmentation with risk, the death rate is larger 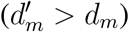 than implied by the fitness landscape death vector **d** (see Eq. (2)).

The equation system (3) is linear with respect to *x*_*i*_. Thus, it can be written in a form of matrix differential equation

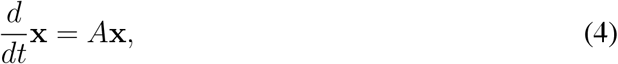

where x = (*x*_1_, *x*_2_, · · ·, *x*_*m*−1_)^*T*^, and matrix *A* is

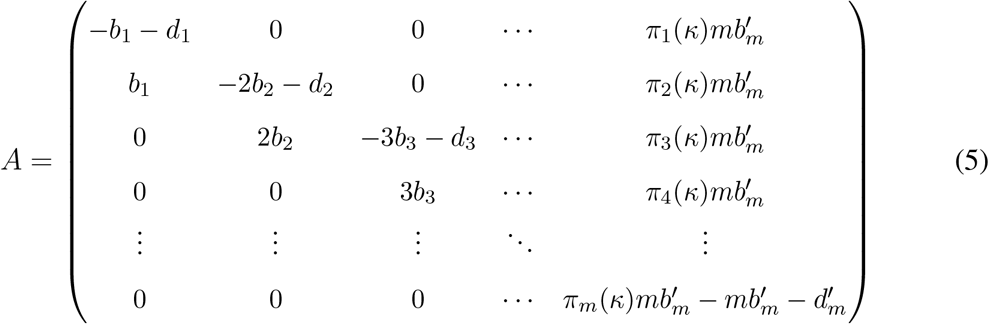

In the long run, the solution of Eq. (4) converges to that of an exponentially growing population with a stable distribution, i.e.,

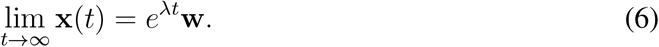

The leading eigenvalue λ gives the total population growth rate, and its associated right eigenvector **w** = (*w*_1_,…, *w*_*m*_) gives the stable distribution of group sizes. In the long term, the fraction of groups of size *i* in the population is proportional to *w*_*i*_. The leading eigenvalue determines the evolutionary success of a population: In the competition of two populations utilizing different life cycles (and hence different λ), the one with larger growth rate will outcompete the other one. Thus, natural selection would promote the life cycle that provides the largest λ. We call this the evolutionary optimal life cycle.

To find the evolutionary optimal life cycle, it is necessary to find values of λ for all life cycles of interest. The leading eigenvalue λ is given by the largest solution of the characteristic equation

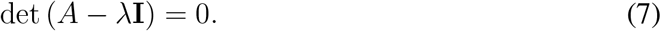

For a given deterministic life cycle associated to fragmentation at size *m* according to the partition *κ*, the characteristic equation (7) reduces to (see Appendix A.2 for a derivation)

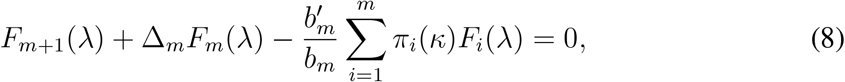

where

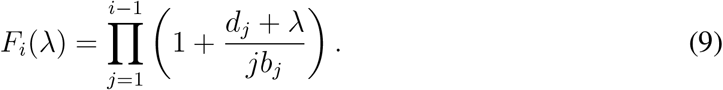

The parameter

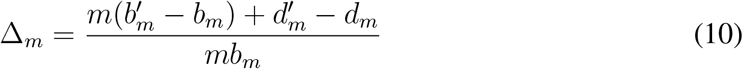

characterises how costly fragmentation is in terms of risks and delays. In the absence of any costs, we have Δ_*m*_ = 0. Eq. (8) is a polynomial equation of degree *m*. In general, we have to solve this equation numerically.

### 2.5 Random fitness landscapes

We now numerically investigate the distribution of optimal life cycles on two sets of fitness landscapes: random fitness landscapes and random detrimental fitness landscapes, which strongly disfavour groups. Both sets are explored by 10000 fitness landscapes generated only once and then used to assess all three scenarios: delay, risk, and loss. Within the scope of this study, we are interested in proportion of fitness landscapes promoting each of the classes of life cycles. The amount of collected data provides the relative accuracy about 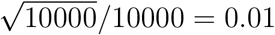, which is enough for our purposes.

In the set of random fitness landscapes, each element of the birth and death rates vector (b and d) was sampled independently from the uniform distribution *U*(0,1).

In the set of random detrimental fitness landscapes, for each landscape, we initially sampled two sequences of *n* = 19 random numbers, each using the uniform distribution *U*(0,1). Then, the first sequence has been sorted in descending order to form the vector of the birth rates b and the second sequence has been sorted in ascending order to form the vector of death rates d. Thus, in all detrimental fitness landscape, the values of birth rates monotonically decreased with the group size, while the values of death rates monotonically increased. Therefore, one could assume that life cycles that fragment at large group sizes only are strongly disfavoured.

## 3 Results

### 3.1 Some life cycles cannot be evolutionary optimal under any fitness landscape

To find which life cycles can evolve for costly fragmentation we consider a large population of groups that can grow without constraint (see Section 2.4). The growth of any group is limited by the maximal group size *n* =19. This leads to 2693 possible life cycles, one for each non-trivial partition of all integers not exceeding 20. The growth rate of a population with any given life cycle can be computed by solving Eq. (8). For each combination of the fitness landscape (Section 2.1) and the fragmentation cost (Section 2.3), one of the 2693 life cycles provides the largest growth rate and, thus, is evolutionary optimal.

For any fitness landscape, it is possible to find a life cycle which is evolutionary optimal under this fitness landscape. However, the opposite is not true: for some life cycles, it is impossible to find any fitness landscape under which it is evolutionary optimal. We label these life cycles “forbidden life cycles”. Consequently, we call a life cycle that is evolutionary optimal under some fitness landscape “allowed life cycle”.

It can be shown analytically that all three scenarios of the fragmentation cost (delay, risk and loss) lead to the same condition for a life cycle to be forbidden: the life cycle determined by the partition *κ* is forbidden if two different subsets of offspring with equal combined sizes exist, i.e. if two partitions *τ*_1_ and *τ*_2_ exist such as:

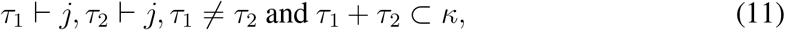

For any fitness landscape and any fragmentation cost scenario, the life cycle employing such a partition is dominated by one of two life cycles in which one of the subsets occurs twice, while other one is not present, see Appendix A.3 for the proof.

The simplest example of the forbidden life cycle is the partition 2+1+1, which has two different offspring subsets: 2 and 1+1, both having the same combined size 2. It is always dominated either by a life cycle with partition 2+2 (subset 2 occurs twice) or by a life cycle with partition 1+1+1+1 (subset 1+1 occurs twice), see Fig. 2a for more examples. The proportion of forbidden life cycles rapidly increases with the partition sum (see black bars on Fig. 2b). Individually assessing each of considered 2693 partitions computationally, we found only 687 partitions corresponding to allowed life cycles (this is about a quarter of the total number).

The total amount of allowed life cycles is still too large to track each of them individually. Therefore, a classification is necessary. We focus on three significant subsets: binary fragmentation, equal split and seeding, see also Fig. 2a. Binary fragmentation partitions have the form *κ* = *a* + *b*. Examples of binary partitions are 2+2 and 7+1. Binary fragmentation cover all scenarios where the parent group divides in two parts. Among the non-binary fragmentation modes, we distinguish equal split and seeding partitions. Equal split partitions have the form *κ* = *a* + ··· + *a* + *b* such that *a* > *b* ≥ 0 and have more than two parts. Examples of equal splits are 1+1+1 and 3+3+3+2. Equal splits represent scenarios, where cells are evenly distributed among multiple offspring groups (plus a single smaller remainder group, if needed). Seeding partitions have the form *κ* = *a* + *b* + ··· + *b* such that *a* > *b* + ··· + *b*. Examples of seeding are 3+1+1 and 7+2+2+2. Distinguishing the seeding fragmentation modes is inspired by seeding dispersal exhibited by biofilms, where a small portion of cells leaves the parent group in an act of fragmentation.

All binary, equal split and seeding partitions are associated to allowed life cycles. However, not every allowed partition belongs to either of these three subsets. For instance, the allowed partitions 4+2+1 and 5+4+4 do not belong to any of these classes. The proportion of binary, equal split and seeding partitions among all allowed partitions decreases with the partition sum, see Fig. 2b). For a system where groups may grow up to *n* = 19, there are 100 binary partitions, 90 equal split partitions, 110 seeding partitions and 387 other allowed partitions, which do not belong to either of these three classes.

**Figure 2:**
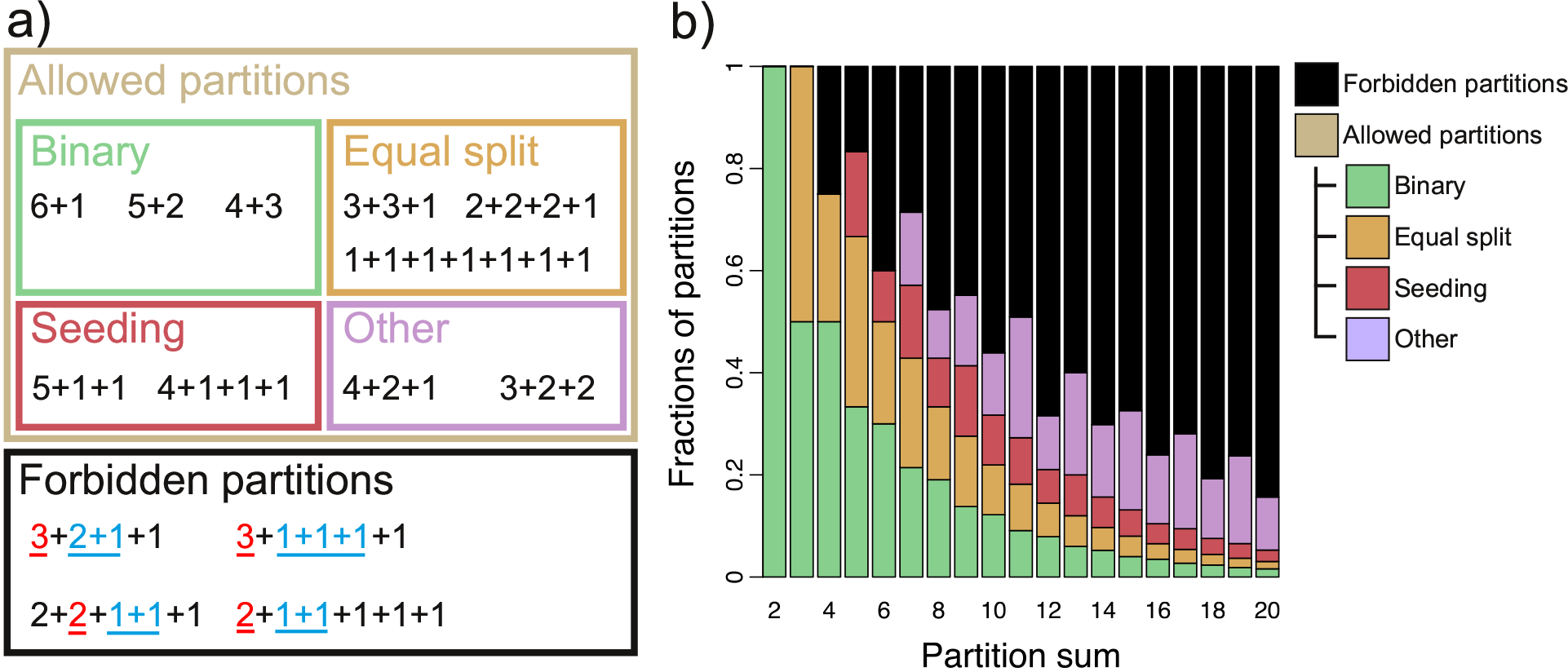
Forbidden and allowed partitions. *(a)* Allowed and forbidden partitions of 7. Allowed partitions are further broken into binary, equal, seeding, and other classes, according to the definitions in the main text. For each of forbidden partitions, a couple of different subsets of parts with the same sum are underlined (see Eq. (11)). *(b)* Proportion of forbidden and allowed partitions as a function of the partition sum. For partition sums 2 and 3 all partitions are allowed, starting from 4 some partitions are forbidden (for partition sum 4, it is 2+1+1). The proportion of forbidden partition grows rapidly with the partition sum. Among allowed partitions, the proportions of binary, equal split and seeding classes rapidly declines, consequently the other partitions constitute the majority of allowed fragmentation modes at large partition sums.

### 3.2 Evolutionary optimal life cycles under random fitness landscapes

The previous section introduced the range of potentially optimal life cycles, but it did not give any insight about interconnection between life cycles and fitness landscapes. Some life cycles may be evolutionary optimal under a larger set of fitness landscapes than others. To study the distribution of optimal life cycles for costly fragmentation, we generated a large set of 10000 random fitness landscapes (see section 2.5). For each fitness landscape from this set, we numerically computed the optimal life cycle independently for each of three scenarios of the fragmentation cost (delay, risk, or loss) under a range of cost values (*T*, *R*, or *L*, respectively).

#### 3.2.1 The average maturity size and the number of produced offspring increase with the increase in fragmentation cost

The average maturity size *m* at which fragmentation occurs and the average size of offspring groups are presented in Fig. 3 a-c. For all three scenarios of the costly fragmentation, the maturity size increases with the cost (*T*, *R*, or *L*). For our choice of *n* = 19, the average maturity size approaches 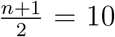 with an increase in fragmentation delay (*T*) and the variation approaches 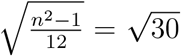 (see Fig. 3a), because the distribution of maturity sizes approaches a uniform distribution, see Appendix A.4. For fragmentation with risk, the average maturity size steadily grows with risk (*R*), while the variation of maturity sizes slowly decreases (see Fig. 3b). For fragmentation with losses, the average maturity size steadily increases with cell loss (*L*) and the variance decreases. At *L* = *n* − 1 = 18 the maturity size is aways *m* = *n* = 19, see Fig. 3c.

Also, the number of offspring increases with the cost. For costless fragmentation, the optimal life cycle always produces exactly two offspring groups. With increasing costs, life cycles with fragmentation into multiple parts become optimal, and consequently, the number of produced offspring increases. For fragmentation with delay, the average size of offspring does not change significantly with delay (*T*), see Fig. 3a. For fragmentation with risk, the average size of offspring decreases with risk (*R*), see Appendix A.5. Combined with the increase in the maturity size, this leads to an increase in the number of offspring produced at the fragmentation event. For fragmentation with loss, the size of offspring monotonically decreases with loss (*L*) and therefore, the offspring number initially increases with loss. However, the number of offspring declines at large *L*, because this number cannot exceed the number of surviving cells, which is limited by *n* − *L* + 1. In our model the number of produced offspring returns to 2 at *L* = 18.

**Figure 3:**
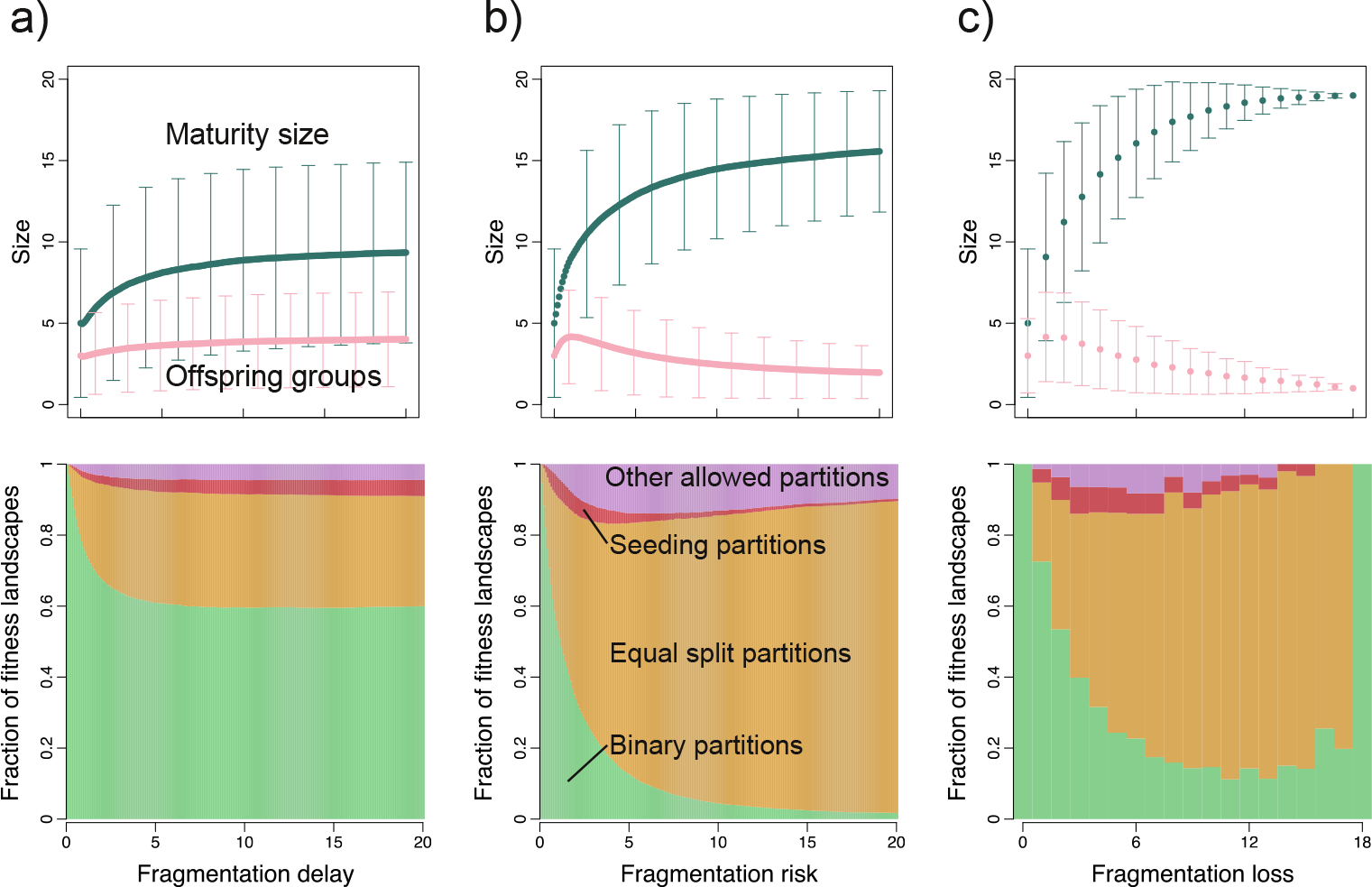
Optimal life cycles for costly fragmentation. The top panels present maturity and average offspring sizes in optimal life cycles as a function of the fragmentation cost for a) fragmentation with delay, b) fragmentation with risk and c) fragmentation with cell loss, respectively. Points depict the average value, error bars represent one standard deviation. The bottom panels show the fractions of each of binary fragmentation, equal split, seeding, and other allowed partitions as functions of fragmentation cost for the same scenarios of fragmentation cost. While the binary and equal split transitions constitute relatively small portion of available partitions, the corresponding life cycles have high probability to be evolutionary optimal. The increase in fragmentation loss reduces the amount of available life cycles, especially at large *L*. Thus, the fraction of life cycles classes at the panel c) does not change smoothly with the fragmentation loss.

#### 3.2.2 Equal split and binary fragmentation life cycles are overrepresented for random fitness landscapes

The proportions of different classes of partitions among optimal life cycles change with the fragmentation cost (*T*, *R*, or *L*), see Fig. 3 a-c.

If reproduction is costless, only binary partitions can be evolutionary optimal [Pichugin et al., 2017]. At low reproduction costs, binary partitions remain the most abundant class under any scenario of cost implementation. With an increase in costs, the fraction of fitness landscapes promoting binary fragmentation declines. For reproduction with delay, this fraction stabilizes at about 60% (see Fig. 3a), while for reproduction with risk, it falls below 5% (see Fig. 3b). For fragmentation with loss, the binary fragmentation increase in abundance up to *L* ≈ 15 on, see Fig. 3c. This is connected to the decrease in the number of available partitions once the fragmentation loss become compatible with the maximal available group size *L* ~ *n* (such as at *L* = 18, the only possible partition is 1+1, which is a binary one).

Equal split fragmentations constitute another major class of observed reproduction modes. For reproduction with risk and with (moderate) losses, equal splits are evolutionary optimal for the vast majority of fitness landscapes. For reproduction with delay, equal splits are the second most abundant class of optimal life cycles. Equal splits are promoted by natural selection, because they maximize the number of offspring groups per act of fragmentation and thus share the cost among the largest number of offspring groups.

Seeding and other fragmentation modes contribute only a small portion of optimal life cycles in all three scenarios of reproduction cost. For reproduction with delay and loss, both these classes are evolutionary optimal at roughly the same proportions of fitness landscapes (~ 5%). Given that there is a much smaller number of seedings than other partitions, see Fig. 2b, seeding partitions are less suppressed by fragmentation with delay and loss than other partitions. For reproduction with risk, seeding partitions are much less abundant than other partitions.

### 3.3 Fragmentation cost can drive the formation of multicellular groups

Multicellular groups evolve when the existence of cells in a group provides some benefit, expressed for example in a form of better resource acquisition or protection from external threats. However, for costly group fragmentation, even when existence in groups is detrimental to cells comprising them, formation of multicellular groups may be evolutionary beneficial: We have constructed a set of 10000 random detrimental fitness landscapes (see Section 2.5). For each of them, the death rate increases monotonically with the size of group, while the birth rate monotonically decreases with the group size. For costless fragmentation, the optimal life cycle for all detrimental fitness landscapes is unicellular, i.e. uses the partition 1 + 1. With the increase in the value of the fragmentation cost (*T*, *L* or *R*), other - multicellular - life cycles become optimal (see Fig. 4). For all detrimental fitness landscapes and all scenarios of the fragmentation cost, all observed optimal life cycles are equal splits in the form 1 + 1 + · · · + 1 (see Fig. 4a-c). The intuition behind this behaviour is that a solitary cell is the most effective state available to the population under the detrimental fitness landscapes, since solitary cells have the largest growth and the lowest death rate among all possible groups sizes.

## 4 Discussion

A key factor considered in the present study is the cost of reproduction - an act of making offspring results in less net biomass than the growth without reproduction. How much is it the case for the natural populations? A number of evidences from observations and experimental studies shows that reproduction can be indeed costly. Such a costs come in different forms. For instance, consider streptococcus bacteria, which naturally forms cell chains held together by cell walls. To fragment, these cell walls must be broken and the process of unchaining requires the expression of autolysin [Lominski et al., 1958, Shaikh and Stewart-Tull, 1975, Mou et al., 1976]. Autolytic-defective mutants unable to fragment and form long chains [Soper and Winter, 1973, Shungu et al., 1979]. The necessary investment of resources into autolysin production constitutes the cost of group fragmentation in this case (represented by the scenario of fragmentation with delay in our model).

**Figure 4:**
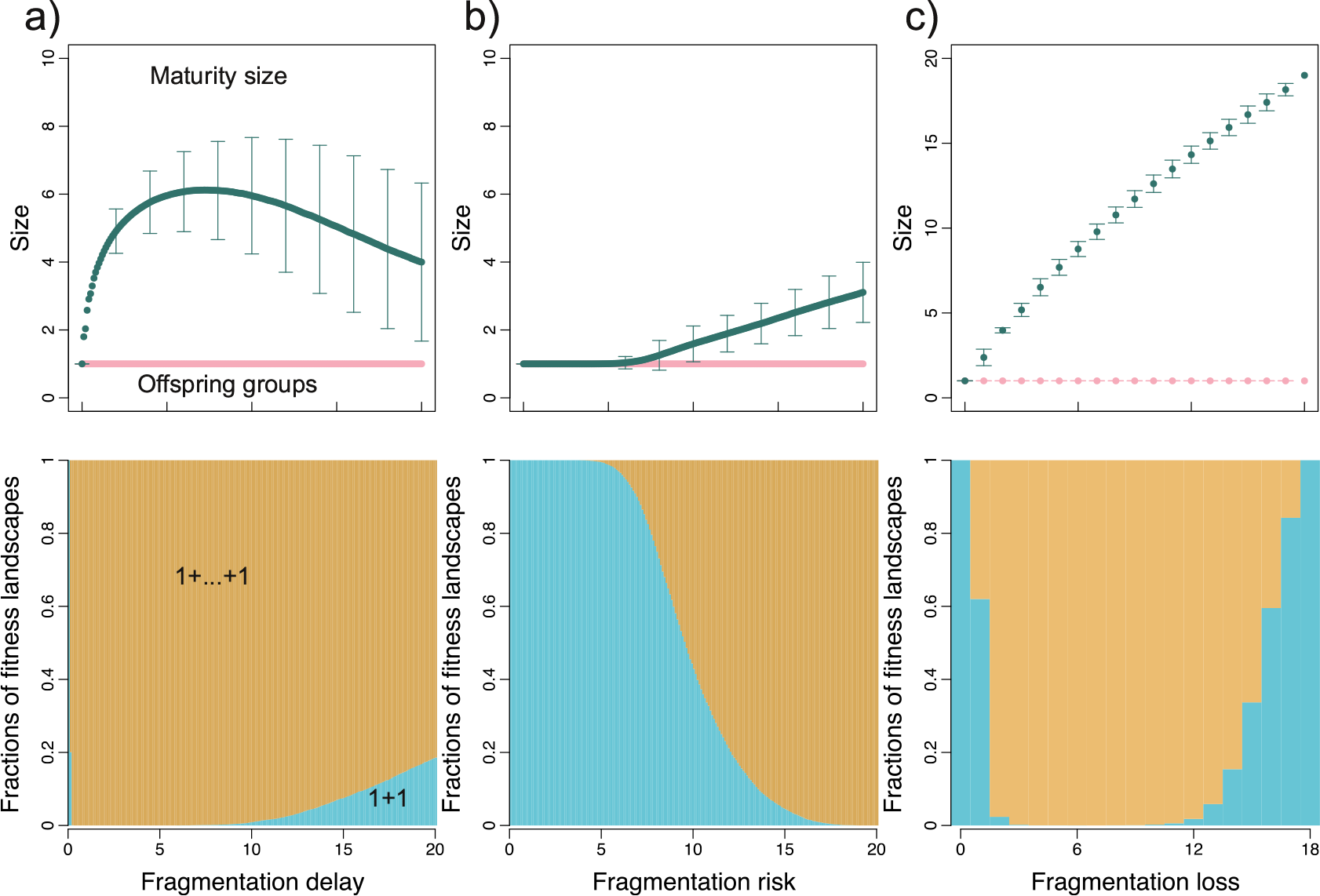
Fragmentation cost can drive the evolution of multicellular groups in detrimental fitness landscapes. The top panels show the average size of the parental and offspring groups in optimal life cycles as a function of fragmentation cost for a) fragmentation with delay, b) fragmentation with risk and c) fragmentation with cell loss, respectively. Points show the average value, error bars represent one standard deviation. The set of fitness landscapes is given by monotonic random sequences (see main text). The size of offspring groups is strictly one, which means that all observed equal split fragmentations had all offspring being independent cells. The bottom panels shows the fractions of unicellular (1+1) and multicellular modes of fragmentation. At no cost, all detrimental fitness landscapes promote the unicellular life cycles. For fragmentation with delay, the fraction of unicellular life cycles rapidly decreases and approaches zero at *T* = 0.3. However, starting from *T* ≈ 5.9 unicellular life cycles become sometimes optimal again. For fragmentation with risk, multicellular life cycles are not observed below *R* ≈ 3.9. Nevertheless, by *R* ≈ 18.9 under all fitness landscapes, the optimal life cycles become multicellular. For fragmentation with loss, the partition 1 + 1 corresponds to the fragmentation at the minimal possible size. The proportion of fitness landscapes promoting the partition 1+1 decreases rapidly. However, the number of available fragmentation partitions decreases with *L* such as at *L* = 18 the only possible partition is 1+1.

Another example is seeding dispersal in bacterial biofilms. Here, the biofilm composed of mostly sessile cells develop cavities filled with motile cells, who are then released into the environment [Webb et al., 2003a, McDougald et al., 2012, Claessen et al., 2014]. To develop cavities and motile cells, the biofilm changes its structure [Purevdorj-Gage et al., 2005], which inevitable bears an investment costs. Moreover, to free up the space for motile cells and provide nutrients for the differentiation, cells in the cavity die [Tolker-Nielsen et al., 2000, Webb et al., 2003b]. Therefore seeding in biofilms is related not to one but to two scenarios of reproduction cost considered in our model.

A unique mechanism of group fragmentation has been developed by *S. cerevisiae* colonies in experimental evolution studies [Ratcliff et al., 2013, 2014]. There an initially unicellular budding yeast was subjected to the selection regime favouring formation of cell clusters. Evolved clusters have a tree-like structure. To facilitate a fragmentation, a single cell in the centre of the tree dies, thus, the integrity of the tree cannot be maintained and eventually the colony breaks into several smaller parts. The death of cell is fragmentation cost in this example. While not being a natural world example, this organism shows that in the need of developing an efficient group fragmentation mode, evolution readily accepts the incurring reproduction costs.

Comparing our results with the case of costless fragmentation considered in Pichugin et al. [2017] suggests that the evolution of life cycles involving fragmentation into multiple parts may be linked with costly group reproduction. Whether this is an actual driving force of evolution in natural populations is an open experimental question. Nevertheless, we can consider known cases of fragmentation into multiple parts and assess whether a group reproduction is associated with any costs.

The first example is the bacterium *Metabacterium polyspora*, inhabiting the gastrointestinal tract of guinea pig. The life cycle of this bacterium involves repeatable passages through the tract of multiple hosts. In order to survive such a process, multiple endospores are produced within a single cell [Angert and Losick, 1998], see Fig. 5a). Up to nine endospores can be formed in a single bacterial cell, which make this life cycle a clear example of a fragmentation into multiple parts. The most apparent cost of reproduction in *M. polyspora* is that the maternal cell is discarded after the release of endospores. Moreover, the formation of endospores in bacteria is significantly different from the normal binary cell division, since the resulting object must survive through much higher stress than the parent cell [Nicholson et al., 2000]. Thus, in addition to the normal machinery involved in DNA replication and cell division, a number of additional processes are involved in production and maturation of the endospore (reviewed in [Angert, 2005]). These processes contribute additional costs of the reproductive event.

Another example is a group of segmented filamentous bacteria [Davis and Savage, 1974], where colonies release two independent cells that grow into new colonies. This reproduction mode can be described by the partition *x* +1 + 1, i.e. it corresponds to the seeding class. The colony of segmented filamentous bacteria originates as a single holdfast-bearing cell, which is capable to attach to the host epithelium. Once this cell settles down, it begins to grow and divide, forming the colony. Since the epithelium is repeatedly renewed tissue, colonies have to give rise to new colonies. This requires production of new holdfast-bearing cells. These cells emerge in a process somewhat similar to the production of endospores - asymmetric division with consequent engulfment of a smaller daughter cell by the larger one. Notably, once the new holdfast-bearing cells have matured, the cell containing them undergoes lysis in order to release them into the gastrointestinal tract, see Fig. 5b). Thus, these organisms pay a similar cost of reproduction.

**Figure 5:**
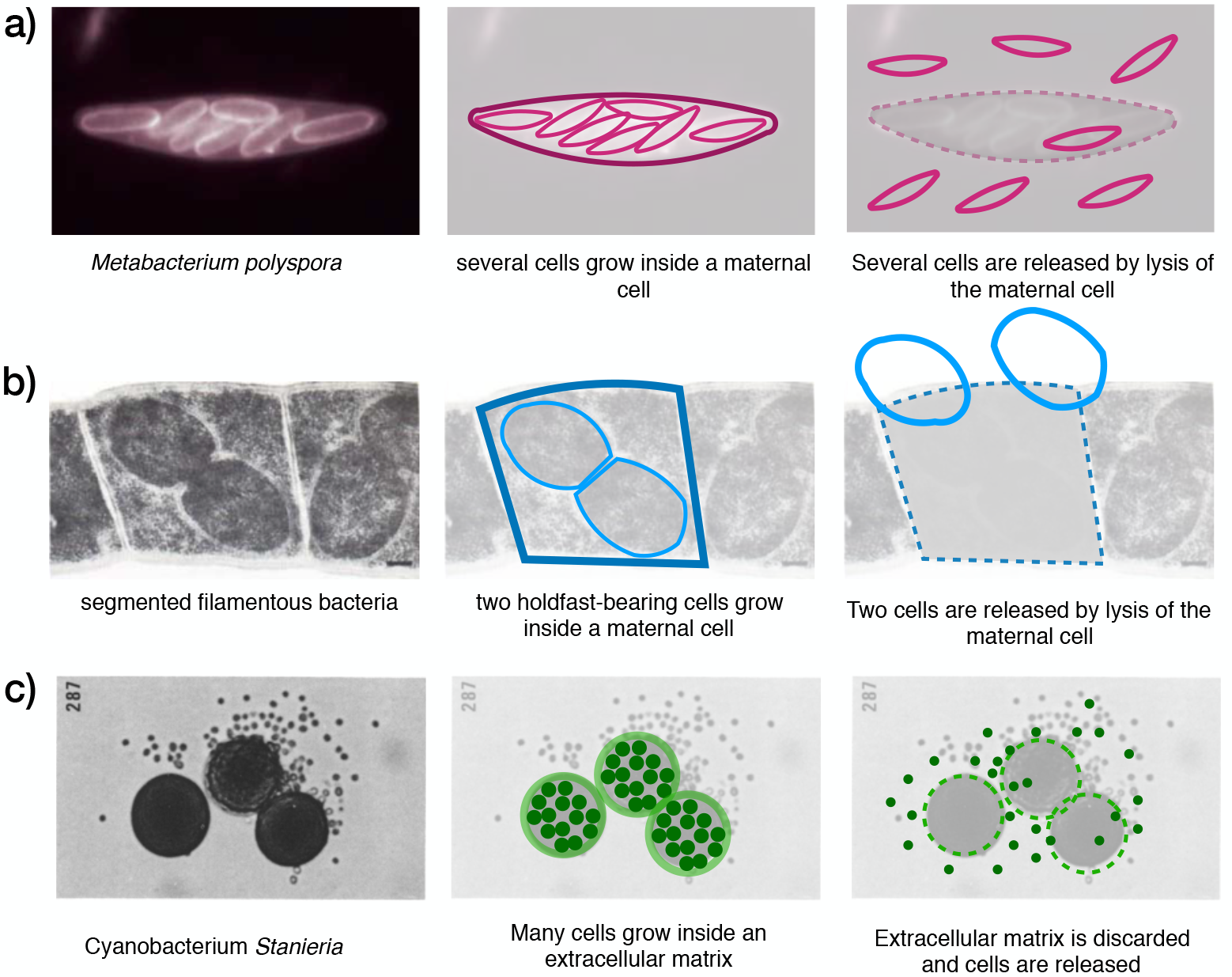
Examples of multiple fragmentation in nature and their interpretation by means of our model. *a) M. polyspora* grows multiple endopsora, released after the maternal cell lysis (picture adopted from [Angert and Losick, 1998]). From the viewpoint of our approach, a group of size *x* + 1 loses one cell and fragments into *x* groups of one cell each. *b)* segmented filamentous bacteria grows two holdfast-bearing cells inside a maternal cell. These cells are released in the result of the maternal cell lysis (picture adopted from [Davis and Savage, 1974]). From the viewpoint of our approach, a group of size *x* + 2 loses one cell and fragments according to the partition (*x* − 1) + 1 + 1. *c)* genus *Stanieria* grows multiple cells within a single body of extracellular matrix. These cells are released simultaneously upon the break of the matrix (Picture adopted from [Waterbury and Stanier, 1978]). From the viewpoint of our approach, a group of size *x* fragments into *x* groups of one cell each, loosing the extracellular matrix, which production required a prior investment of resources.

An example for an organism with fragmentation cost in a form other than cell loss is the *Stanieria* genus of cyanobacteria. These organisms are born as independent cells. In the course of their life, these cyanobacteria continuously produce an extracellular matrix, which helps the organism to attach to solid surfaces. Shortly before the reproductive event, the cells undergo a rapid succession of fissions, producing between 4 and 1000 cells. Then, the extracellular matrix gets broken, releasing multiple offspring at once [Waterbury and Stanier, 1978], see Fig. 5c. In this case, the fragmentation cost comes in the form of the lost extracellular matrix, which protected and sustained the parent organism, but is not transferred to the offspring cells. The production of the extracellular matrix is distributed across the whole lifespan of the organism, therefore, this scenario lays outside of the scope of the current model, where the cost is assumed to be paid at the last step of the organism’s life. Nevertheless, the combination of multiple fragmentation in *Stanieria* and the apparent costs of the reproduction qualitatively support our hypothesis that fragmentation costs can drive life cycle evolution.

Other notable examples of multiple fragmentation, which are even further away from our model are algae *Gonium pectorale* and slime molds. *G. pectorale* also undergoes sexual reproduction, which violates the assumption of asexual reproduction in our model. Slime molds colonies are formed by aggregation of cells and not by the growth of previous member of the colony. Still, both organisms exhibit fragmentation into multiple parts and significant fragmentation costs. *G. pectorale* spends the majority of its life cycle in a form of 16-cell colony. At the fragmentation, the colony dissolves into 16 independent cells, which originate new colonies [Stein, 1958]. Since the maturity size for *G. pectorale* is 16 cells, but the fragmentation does not immediately follow the moment of the reaching this size, this organism has an explicit delay of fragmentation. Slime molds, which are popular model organisms in studies on the evolution of cooperation, form a slime composed of multiple cells. The slime further differentiates into fruiting body containing multiple spores and stalk needed to provide some height to the fruiting body, so spores can be distributed across larger territory [Bonner, 1959]. Cells in the stalk die without contributing to the spores, thus the stalk is the cost of the fragmentation. Both organisms, support the hypothesis as well, but only on a conceptual level.

Another aspect of our study is that all three considered scenarios of the fragmentation cost share the same set of potentially optimal life cycles. For fragmentation with delay, risk and cell loss, only these life cycles, which partitions do not contain two different partitions with the same sum, can be evolutionary optimal. Given the difference between the ways how the considered costs affect the life of a single organism in a population, this result is striking.

For costless fragmentation and fragmentation with proportional costs, only binary fragmentation can be evolutionary optimal [Pichugin et al., 2017], which vastly reduces the number of possible life cycles. For instance, if the group size limited by *n* = 19, there are only 99 binary fragmentations which can evolve for costless fragmentation. The introduction of the fixed fragmentation cost expands the space of optimal life cycles. For costly fragmentation, the number of potentially optimal life cycles is almost 7 times larger: 687.

Among all potentially optimal life cycles, we discriminate two special classes: binary fragmentation and equal split. They constitute only a small fraction of all allowed life cycles, see Fig. 2b. However, these two narrow classes of fragmentation modes are evolutionary optimal under majority of random fitness landscapes for all three scenarios of the fragmentation cost, see Fig. 3. Among the natural bacterial populations and simple eukaryotic species, binary fission is the dominant mode of reproduction (see [Angert, 2005]). The majority of species, which utilize the fragmentation into more than two parts, do it by fission in multiple unicellular propagules, as discussed above. A notable exception is the reproduction mode of segmented filamentous bacteria [Davis and Savage, 1974] (see above). Thus, binary fragmentation and equal split are not only promoted by our model, but also relatively widespread in nature.

The evolution of groups from unicellular ancestors is often considered to be driven by some ongoing benefits provided by the group membership such as better protection [Stanley, 1973], access to novel resources[Rainey and Travisano, 1998] and the opportunity to cooperate (reviewed in [Kaiser, 2001] and in [Grosberg and Strassmann, 2007]). In our work we have shown that such ongoing benefits of being in a group are not a necessary condition for the evolution of groups. Another, previously overlooked factor capable to drive the evolution of groups is the cost regularly paid at each reproduction event. The impact of the reproduction cost is strong enough that it may promote formation of multicelluar groups even if the group living put cells in disadvantage comparing with solitary existence. Two factors contribute to this effect. First, the growth to larger size takes more time and thus makes reproduction less frequent, so the cost per time unit is smaller. Second, larger group size at fragmentation makes it possible to share the burden of reproduction cost among more units. This reduction of the impact of the reproduction cost is previously overlooked factor, which promotes the formation of multicellular groups.

Given the fascinating diversity of biological life cycles observed even in simple organisms, it seems daunting to use theoretical models to understand their features. However, our approach shows that even simple models can capture key aspects of this process and produce results for a whole variety of life cycles. At the same time, these models point towards fragmentation costs as potential drivers of this diversity.

## 5 Acknowledgements

We are grateful to David Rogers and Philippe Remigi for fruitful discussions and biological insights.

## A Appendix

## A.1 Only deterministic fragmentation modes can be evolutionary optimal under any fitness landscape

Following Pichugin et al. [2017], the state of the population can be described by the vector x, where *x*_*i*_ denotes the abundance of groups of size *i*. All processes changing the state vector x - birth, death and fragmentation - occur with a constant rate. Thus, the dynamics of the population state can be described by a set of linear differential equations or, equivalently, by a matrix differential equation

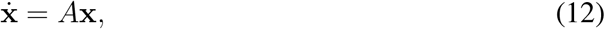

where *A* is a projection matrix defined by demographics of the population [Caswell, 2001]. An element *a*_*i,j*_ of the projection matrix describes the rate of change of the number of groups of size *i* caused by processes occurring with groups of size *j*.

To construct the projection matrix elements, consider groups of a certain size *j*. We denote by *q*_*j,κ*_ the probability that upon the growth from size *j* to *j* + 1, the group will fragment by a partition *κ* ⊢ *j*′ ≤ *j* + 1 (where the “≤” indicates that cells can be lost upon fragmentation). Among these partitions we distinguish the trivial partition of *j* + 1 that corresponds to the growth without fragmentation; we denote this by *q*_*j*,(*j*+1)_. The combined probability of all outcomes is equal to one:

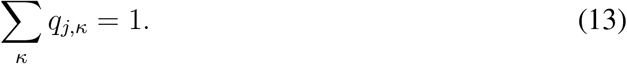

For deterministic life cycles, only one partition occurs in all groups in a population. Thus, for group sizes *j* up to maturity size *m*, the trivial partition occurs with probability one (*q*_*j*,(*j*+1)_ = 1), while all other partitions have zero probability. Once the group grows from the maturity size, a certain non-trivial partition of *j*′ ≤ *m* +1 occurs with probability one. In a stochastic life cycle, more than one partition has non-zero probability at least at one group size. Therefore, the projection matrix is different from Eq. (5).

To show that stochastic life cycles are dominated by deterministic ones, we construct the projection matrix for an arbitrary stochastic life cycle. Groups grow by one cell at a time, thus no process can increase the size of group by more than one unit at once, so *a*_*i,j*_ = 0 for all *i* > *j* + 1. Thus, the projection matrix may contain non-zero elements only in the upper right triangle (emergence of smaller groups during fragmentation), on the main diagonal (fragmentation, growth and death of clusters), and on the first lower subdiagonal (growth of clusters to sizes larger by one cell).

The first lower subdiagonal describes the rate of emergence of new larger groups in a result of group growth without fragmentation. These rates are equal to the product of the basic growth rate and the probability of the group to grow:

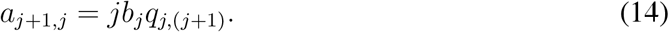

The upper right triangle of the matrix describes the emergence of new groups in a result of fragmentation of larger groups. For a given partition κ and given size of the newborn group i, the rate of production of new groups is equal to the product of the fragmentation rate 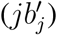, the probability to fragment according to the given partition (*q*_*j,κ*_), and the number of groups of given size produced in the act of fragmentation with this partition (*π*_*i*_(*κ*)). The value of an element *a*_*i,j*_ in the upper left triangle is equal to the sum of rates provided by all partitions available to groups of size *j*:

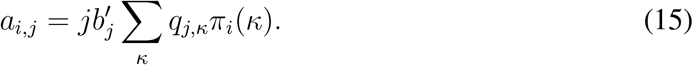

The main diagonal *a*_*i,i*_ describes the changes in groups numbers due to growth and fragmentations as well as the death of groups. The first component of *a*_*i,i*_ is given by the fact that once a group of size *j* grows or fragments, the number of groups of that size decreases. The rates of decrease are equal to *jb*_*j*_*q*_*j*,(*j*+1)_ due to the growth and 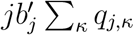 due to the fragmentations. The second component is provided by the fragmentation with partition *κ* = *j* + 1, which produce groups of size equal to the size of parent. This leads to an increase in the number of groups of size *j* at rate 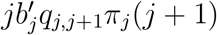, where *π*_1_ (1 + 1) = 2 and *π*_*j*_ (*j* + 1) = 1 if *j* > 1. The last component of *a*_*i,i*_ comes from the death of groups, which leads to a decrease in their number at rate 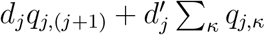, where the first term describes the death rate in the absence of the fragmentation and the second term describes the death rate of fragmenting groups. Combined, the diagonal elements of projection matrix are

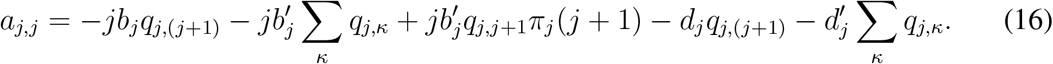

All elements of the projection matrix given by Eqs. (14)–(16) are linear with respect to any probability *q*_*j,κ*_. As shown in Pichugin et al. [2017], in this case the optimal life cycle is always deterministic, independent of the parameter values, such as the fitness landscape and the scenario of the fragmentation cost.

## A.2 Characteristic equation of a deterministic fragmentation mode

Consider a deterministic fragmentation mode in which groups grow up to the maturity size *m* and once the next cell is born, fragment according to a partition *κ* ⊢ *j*′ ≤ *m* + 1. The corresponding projection matrix is an *m* × *m* matrix of the form

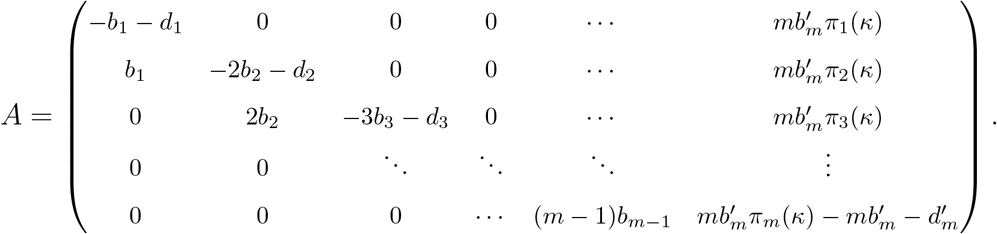

The population growth rate is given by the leading eigenvalue λ_1_ of *A*, i.e., the largest solution of the characteristic equation

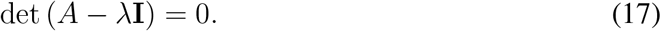

By using a Laplace expansion along the last column of *A* − λ**I**, we can rewrite the left hand side of the above expression (i.e., the characteristic polynomial of *A*) as

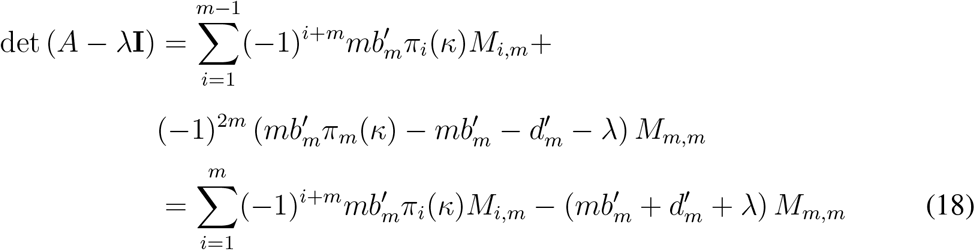

where *M*_*i,m*_ is the (*i, m*) minor of *A* − λ**I**. For all *i* = 1,…, *m*, the minor *M*_*i,m*_ is the determinant of a block diagonal matrix, and hence equal to the product of the determinants of the diagonal blocks. Moreover, each diagonal block is either a lower triangular or an upper triangular matrix, whose determinant is given by the product of the elements in their main diagonals. We can then write

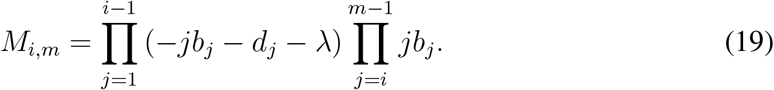

Substituting Eq. (18) and Eq. (19) into Eq. (17) and simplifying, we obtain

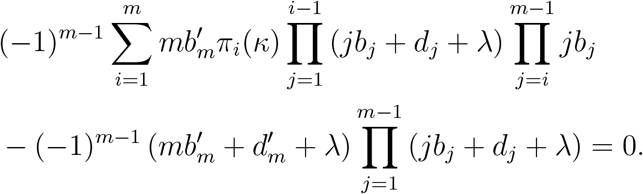

Dividing both sides by

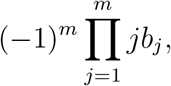

we get

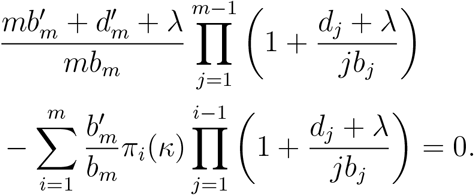

To move the first multiplier with λ into the product, we rewrite it as

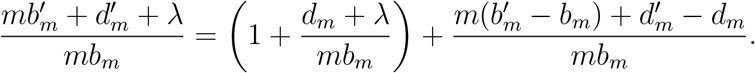

Thus,

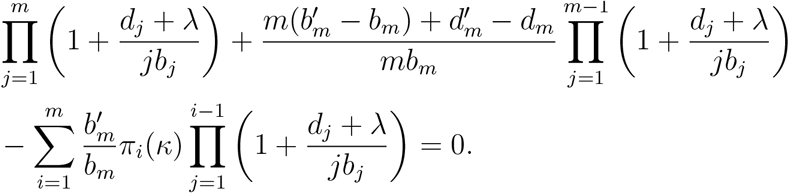

Simplifying this, we finally obtain that the characteristic equation (17) can be written as

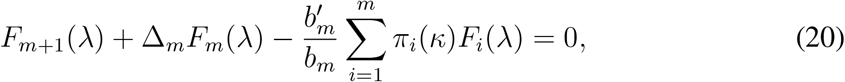

where

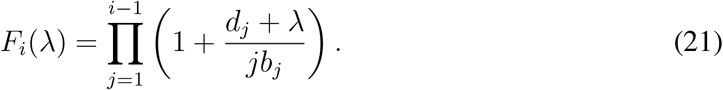

and

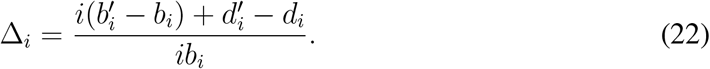

Note that two transformations preserve Eq. (20):

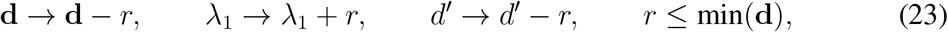

and

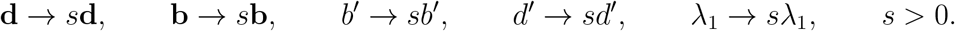

These transformations allow us to set *b*_1_ = 1 and min(**d**) = 0 without loss of generality.

## A.3 Forbidden fragmentation modes

For any fitness landscape, for any combination of the fragmentation delay, risk and fixed loss, the fragmentation mode having two different subsets of offspring with the same combined size is dominated. To prove this, we use approach similar to one used in Appendix E in [Pichugin et al., 2017]. Consider positive integers *m*, *j*, *k* such that *m* + 1 ≥ 2*j* + *k*, two partitions *τ*_1_ ⊢ *j* and *τ*_2_ ⊢ *j* such that *τ*_1_ ≠ *τ*_2_, and an arbitrary partition *ϕ* ⊢ *k*, and the following three deterministic fragmentation modes:

1. *κ*_1_ = *τ*_1_ + *τ*_2_ + *ϕ* ⊢ 2*j* + *k* ≤ *m* +1, whereby a complex fragments upon growth from size *m* into a number of offspring given by partitions *τ*_1_, *τ*_2_, and *ϕ*.
2. *κ*_2_ = *τ*_1_ + *τ*_1_ + *ϕ* ⊢ 2*j* + *k* ≤ *m* + 1, whereby a complex fragments upon growth from size *m* into a number of offspring given by two partitions *τ*_1_ and one partition *ϕ*.
3. *κ*_3_ = *τ*_2_ + *τ*_2_ + *ϕ* ⊢ 2*j* + *k* ≤ *m* + 1, whereby a complex fragments upon growth from size *m* into a number of offspring given by two partitions *τ*_2_ and one partition *ϕ*.

Denoting by λ(*κ*_*i*_) the leading eigenvalue of the projection matrix induced by fragmentation mode *κ*_*i*_, we can show that, for any fitness landscape, either λ(*κ*_1_) ≤ λ(*κ*_2_) or λ(*κ*_1_) ≤ λ(*κ*_3_) holds. This means that a fragmentation mode with two different subsets of offspring with the same combined size is dominated by a mode where one of these subsets repeats twice, while another one is not present.

To prove the statement above, let us define the polynomial *p*_*i*_(λ) as the left hand side of Eq. (20) with *κ* = *κ*_*i*_, so that λ(*κ*_*i*_) is the largest root of *p*_*i*_(λ). We obtain

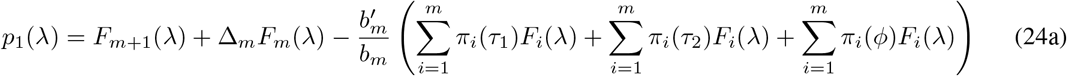

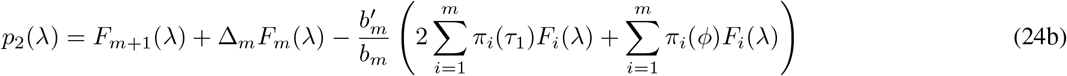

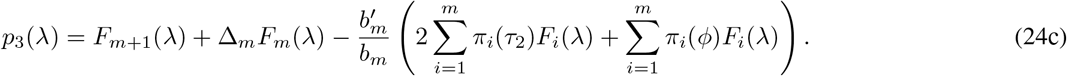

These polynomials satisfy the following two properties. First,

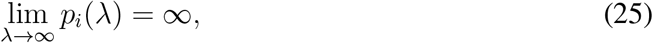

as the leading coefficient of the left hand side of Eq. (20) is given by (*b*_1_ · *b*_2_ · … · *b*_*m*_*m*!)^−1^, which is always positive. Second,

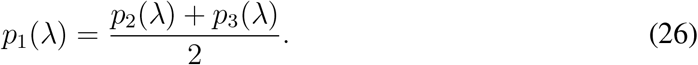

Evaluating Eq. (26) at λ(*κ*_1_), and since λ(*κ*_1_) is a root of *p*_1_(λ), *p*_1_(λ(*κ*_1_)) = 0, it then follows that

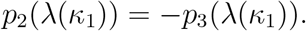

Hence, it must be that only one of the following three scenarios is satisfied: (i) *p*_2_(λ(*κ*_1_)) < 0 < *p*_3_(λ(*κ*_1_)), (ii) *p*_2_(λ(*κ*_1_)) = *p*_3_(λ(*κ*_1_)) = 0, or (iii) *p*_2_(λ(*κ*_1_)) > 0 > *p*_3_(λ(*κ*_1_)). If *p*_2_(λ(*κ*_1_)) < 0 < *p*_3_(λ(*κ*_1_)), and by virtue of Eq. (25) and Bolzano’s theorem (if a continuous function has values of opposite sign inside an interval, then it has a root in that interval), *p*_2_(λ) has a root between λ(*κ*_1_) and ∞. Therefore, λ(*κ*_1_) ≤ λ(*κ*_2_) holds. Likewise, if *p*_2_(λ(*κ*_1_)) > 0 > *p*_3_(λ(*κ*_1_)), then λ(*κ*_1_) ≤ λ(*κ*_3_) holds. Finally, if *p*_2_(λ(*κ*_1_)) = *p*_3_(λ(*κ*_1_)) = 0, then both λ(*κ*_1_) ≤ λ(*κ*_2_) and λ(*κ*_1_) ≤ λ(*κ*_3_) hold. We conclude that either λ(*κ*_1_) ≤ λ(*κ*_2_) or λ(*κ*_1_) ≤ λ(*κ*_3_) must hold.

## A.4 Optimal life cycles under large delay of fragmentation

Consider the deterministic life cycle that follows partition *κ*. Its proliferation rate is given by Eq. (8). Under fragmentation with delay, the birth rate at the fragmentation size changes according to

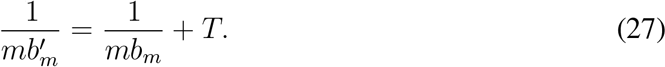

At large delay 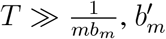 can be approximated as

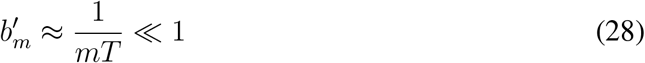

Thus, Δ_*m*_ given by Eq. (22) can be approximated by:

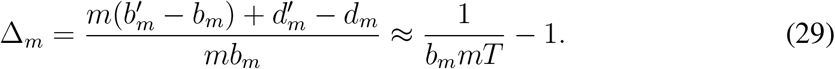

Therefore, Eq. (20) becomes:

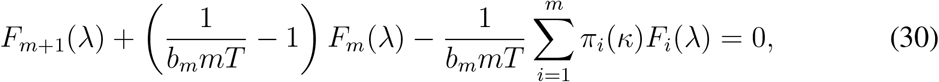

The delay value *T* contribute to this equation only in a form of factor 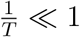. To analyse the solutions of obtained equation, we first discard all terms containing 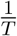 in Eq. (30) and get

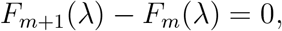

Substituting the expression of *F*_*i*_(λ) from Eq. (21) we get

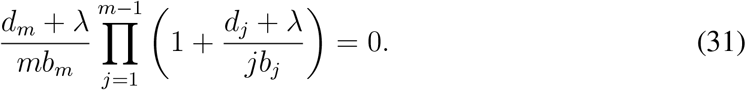

We denote the solutions of this equation as λ^0^. There are *m* solutions of this equation: one solution 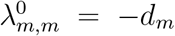 and *m* − 1 solutions in a form 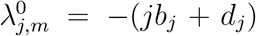, where *j* ∊ {1, 2, · · ·,*m* − 1}. For any solution in a form 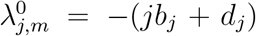, we can find another life cycle fragmenting already at size *j* < *m* for which the solution 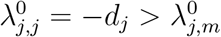 exists. Thus, the proliferation rate of the optimal life cycle must have the form 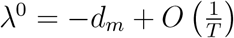. As a consequence, for high fragmentation delay, under the optimal life cycle, group fragments after reaching the most protected state with the minimal *d*_*i*_.

To find which of many fragmentation modes available to the group reproducing at the most protected state is evolutionary optimal, we consider the first order approximation of the growth rate given by

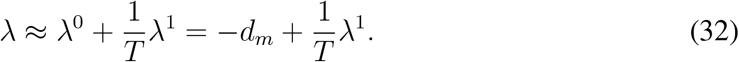

To find λ_1_ we substitute Eq. (32) into Eq. (30),

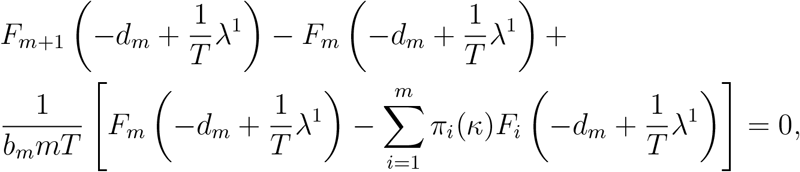

Then we use expressions of *F*_*i*_(λ) from Eq. (21) and discard all terms smaller than 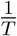

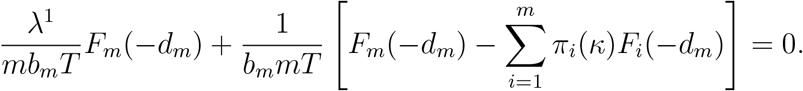

Thus,

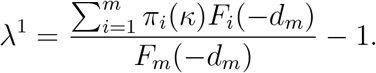

In the optimal life cycle under high delay of fragmentation, groups fragment according to the partition that provides the highest value of λ^1^.

For the special case of the constant death rate, the optimal life cycle can be found explicitly. In this case, the death rate can be set to **d** = 0 (see Eq. (23)), so

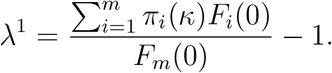

At **d** = 0, *F*_*i*_(0) = 1, so:

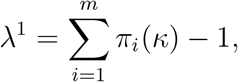

the right hand side of this expression is the number of produced offspring groups minus one. This expression is maximized by the life cycle producing the maximal number of offspring groups, i.e. by the equal split life cycle producing only unicellular propagules.

For the set of random fitness landscapes used in section 3.2, the minimum of *d*_*i*_. is evenly distributed across all considered sizes {1, 2, · · ·, 19}. Thus, at large delay, the size of fragmentation should be evenly distributed as well, which corresponds to average fragmentation size equal to 10 and standard variation of sizes equal to 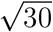. The mean and standard variation of the observed distribution of fragmentation sizes quickly approach these values in our numerical simulations, cf. Fig. 3.

For the set of fitness landscapes detrimental to larger groups used in section 3.3, the minimum of *d*_*i*_ is achieved always at *d*_1_. Therefore, the maturity size for large delay is 1, which corresponds to the unique fragmentation pattern 1 + 1. In our simulations the initial increase in *T* resulted in the gradual decrease of the fraction of fitness landscapes promoting 1 + 1 to zero. However, further increase of *T* make some fitness landscapes promote 1 + 1 again, and above some intermediary value of *T*, the fraction of fitness landscapes promoting 1 + 1 begin to increase, see Fig. 4a.

## A.5 Optimal life cycles under high risk of fragmentation

Consider the deterministic life cycle that follows partition *κ*. It’s proliferation rate is given by Eq. (8). For fragmentation with risk, the death rate at the fragmentation size changes according to

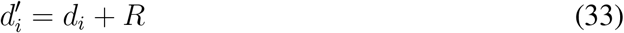

Thus, Δ_m_ given by Eq. (10) becomes

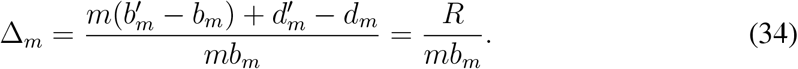

Therefore, Eq. (8) becomes

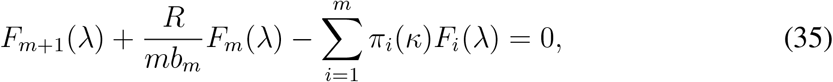

Or, after dividing by *R*,

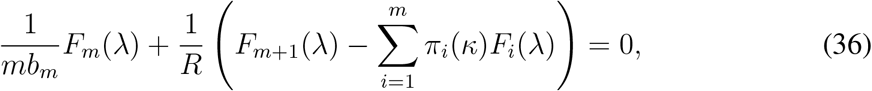

To analyse the solutions of obtained equation, we first discard all terms containing 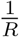 and get

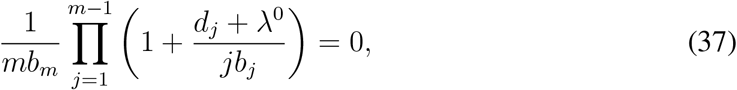

For *m* > 1, this equation has *m* − 1 solutions in a form λ^0^ = −*jb*_*j*_ − *d*_*j*_. Thus, the first approximation of the proliferation rate, given by the maximal root of this equation, is equal to

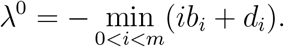

For *m* = 1, this equation has no solution, instead the proliferation rate of the population undergoing 1+1 life cycles (the only life cycle with *m* = 1) is given by

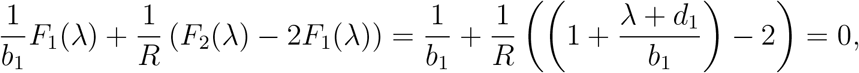

Thus, for *κ* = 1 + 1, the proliferation rate is given by

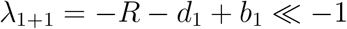

Thus, under high risk of fragmentation, the life cycle with *κ* = 1 + 1 is dominated by any other life cycle. Accordingly, in our simulations, the proportion of unicellular life cycles monotonically decreases with the increase in *R*, see Figs. 3e) and 4. Therefore, according to the approximation, natural selection promotes life cycles with maturity size *m* greater than the critical value *i*\* minimizing expression *ib*_*i*_ + *d*_*i*_.

To distinguish between such life cycles, we consider the first order approximation of the growth rate given by

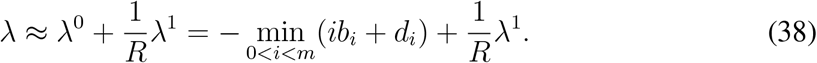

We substitute λ in the form of Eq. (38) into Eq. (36) and discard all terms smaller than 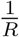

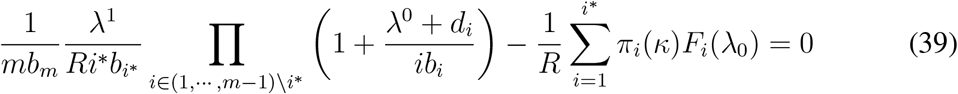

Note, that offspring groups of size larger than *i*\* do not contribute to sum at the end of the expression at the left hand side, because *F*_*i>i\**_(λ^0^) = 0. The term linear with respect to 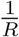 is equal to

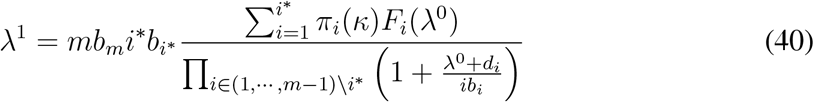

The optimal life cycle maximizes this expression.

For any given maturity size *m*, the life cycle producing more offspring groups with size not exceeding than *i*\* has higher λ^1^. Thus, under the optimal life cycle, the size of offspring cannot be larger than *i*\*.

For the special case where **d** = 0 and *b*_*i*_ does not decrease faster than *i*^−1^, the sequence *ib*_*i*_ + *d*_*i*_ monotonically increases. Hence, *i*\* = 1, so the optimal life cycle is the fragmentation into unicellular propagules.

For the set of random fitness landscapes used in section 3.2, the expression *ib*_*i*_ + *d*_*i*_ tend to grow with *i*, so its minimum *i*\* is more likely to be achieved at small values of *i*. Since, *i*\* establishes an upper limit on the size of offspring groups, our analysis suggests that this size should decrease with *R*.

